# Collaborative metabolic curation of an emerging model marine bacterium, *Alteromonas macleodii* ATCC 27126

**DOI:** 10.1101/2023.12.13.571488

**Authors:** Daniel Sher, Emma E. George, Matthias Wietz, Scott Gifford, Luca Zoccaratto, Osnat Weissberg, Coco Koedooder, Bashir VK Waseem, Marcelo Malisano Barreto Filho, Raul Mireles, Stas Malavin, Michal Liddor Naim, Tal Idan, Vibhaw Shrivastava, Lynne Itelson, Dagan Sade, Alhan Abu Hamoud, Yara Soussan, Noga Barak, Peter Karp, Lisa Moore

## Abstract

Inferring the metabolic capabilities of an organism from its genome is a challenging process, relying on computationally-derived or manually curated metabolic networks. Manual curation can correct mistakes in the draft network and add missing reactions based on the literature, but requires significant expertise and is often the bottleneck for high-quality metabolic reconstructions. Here, we present a synopsis of a community curation workshop for the emerging model marine bacterium *Alteromonas macleodii* ATCC 27126 and its genome database in BioCyc, focusing on pathways for utilizing organic carbon and nitrogen sources. Due to the scarcity of biochemical information or gene knock-outs, the curation process relied primarily on published growth phenotypes and bioinformatic analyses, including comparisons with related *Alteromonas* strains. We report full pathways for the utilization of the algal polysaccharides alginate and pectin in contrast to inconclusive evidence for one carbon metabolism and mixed acid fermentation, in accordance with the lack of growth on methanol and formate. Pathways for amino acid degradation are ubiquitous across *Alteromonas macleodii* strains, yet enzymes in the pathways for the degradation of threonine, tryptophan and tyrosine were not identified. Nucleotide degradation pathways are also partial in ATCC 27126. We postulate that demonstrated growth on nitrate as sole N source proceeds via a nitrate reductase pathway that is a hybrid of known pathways. Our evidence highlights the value of joint and interactive curation efforts, but also shows major knowledge gaps regarding *Alteromonas* metabolism. The manually-curated metabolic reconstruction is available as a “Tier-2” database on BioCyc.

**Importance:** Metabolic reconstructions are vital for the systemic understanding of an organism’s ecology. Here, we report the outcome of a collaborative, interactive curation workshop to build a curated “metabolic encyclopedia” for *Alteromonas macleodii* ATCC 27126, a marine heterotrophic bacterium with widespread occurrence. Curating pathways for polysaccharide degradation, one-carbon metabolism, and others closed major knowledge gaps, and identified further avenues of research. Our study highlights how the combination of bioinformatic, genomic and physiological evidence can be harvested into a detailed metabolic model, but also identifies challenges if little experimental data is available for support. Overall, we show how an interactive get-together by a diverse group of scientists can advance the ecological understanding of emerging model bacteria, with relevance for the entire scientific community.

## Introduction

Metabolism, the complex network of (mostly enzymatic) reactions within and between cells, underlies life on Earth. Reconstructing the metabolic network of an organism based on genomic information remains a fundamental challenge in biology (Fang et al. 2020, Bernstein et al. 2021). For model organisms like *Escherichia coli*, decades of physiological, biochemical, molecular and bioinformatic work have resulted in precise maps of cellular metabolism and its regulation (Fang et al. 2020). These maps and accompanying biological knowledge form an integrated “encyclopedia of the cell”, such as the EcoCyc database, helping to explore cell metabolism and interpret experimental results (Karp et al. 2023). Metabolic reconstructions also serve as basis for quantitative and mechanistic models of cell growth under different conditions (Fang et al. 2020). Curated metabolic databases are available for several medically and biotechnologically-relevant model bacteria (e.g. *Salmonella enterica* (Métris et al. 2017) and *Bacillus subtilis* (Pedreira et al. 2021)) as well as for selected eukaryotic organisms such as *Saccharomyces cerevisiae* (https://yeast.biocyc.org), *Arabidopsis thaliana* (Mueller et al. 2003), and humans (Romero et al. 2005). However, the vast majority of metabolic models for thousands of other organisms are derived purely from automatic pipelines for gene and pathway identification, without manual curation (e.g. (Arkin et al. 2018, Karlsen et al. 2018, Karp et al. 2019)).

Although computational reconstructions are useful starting points for understanding cell metabolism, they are often incomplete or incorrect. For example, they may lack metabolic reactions encoded in the genome that were not identified by the computational pipelines that link genes to reactions and products. Furthermore, entire pathways can be incorrectly predicted (“false positives”) based on the presence of only some associated genes, especially if involved in multiple pathways (Bernstein et al. 2021). Additionally, computational reconstructions lack the supervision of a human curator, who can consider supporting experimental evidence.

Manually curating a metabolic reconstruction, such as the “Tier-2” PGDBs (Pathway/Genome DataBase) available on BioCyc.org (Karp et al. 2019), comprises several stages. The initial metabolic model is computed from a published genome. In BioCyc, this is based on prior genome annotation using Pathway Tools (PTools) software (Karp et al. 2021) and MetaCyc (Caspi et al. 2020) as the reference database for metabolic reactions. Gaps in the draft metabolic network are then filled by suggesting candidate genes (“pathway hole filling”, (Green et al. 2004)). For BioCyc this stage includes also the prediction of transport reactions (Lee et al. 2008) and operons, and imports protein features from UniProtKB (The_UniProt_Consortium 2022) and Protein Database (Burley et al. 2021). Finally, manual curation by one or more experienced curators includes correcting errors, updating gene and protein information, and summarizing the presence and function of enzymes, reactions and pathways based on the literature and experimental evidence. Supporting information includes gene knock-outs or mutants, enzymatic activity assays, transcriptomics and proteomics. The curation process also highlights needs for additional experimental verification of specific pathways. Because the manual curation process takes months to years, the BioCyc collection contains, as of November 2023, only 79 manually-curated (“Tier-1 and Tier-2”) PDGBs, compared to nearly 20,000 purely computationally generated ones (“Tier-3”). Thus, manual curation constitutes a significant bottleneck in consolidating knowledge on cellular metabolism, especially in emerging model organisms with environmental, biotechnological or medical potential, for which resources and data are limited.

One way of facilitating high-quality metabolic reconstructions is the joint effort by a community of researchers, either as a decentralized effort to which curators contribute remotely, or as an in-person curation workshop (e.g. (Stein 2001, Elsik et al. 2006)). During the early days of genome sequencing, such community efforts were relatively common (Stein 2001). Today, such efforts focus on annotating individual genes in eukaryotes, and are facilitated by web portals such as Apollo, Jbrowse, ORCAE, and G-OnRamp (Sterck et al. 2012, Buels et al. 2016, Dunn et al. 2019, Liu et al. 2019). Community annotation also serves to update the Gene Ontology database (Ramsey et al. 2021), and is an exciting way to involve undergraduates in bioinformatic research (e.g. (Hosmani et al. 2019, Jung et al. 2020)).

Building upon the community curation approach, and extending it from individual genes to metabolic pathways, we organized an in-person community curation for the metabolic reconstruction of the marine bacterium *Alteromonas macleodii* ATCC 27126 (herein referred to as ATCC 27126). *Alteromonas macleodii* belongs to the ecologically and physiologically diverse genus *Alteromonas*, which is ubiquitous in tropical and temperate oceans and often abundant on particles (Alonso-Sáez et al. 2007, Roth Rosenberg et al. 2021, Henríquez-Castillo et al. 2022, Wietz et al. 2022). *Alteromonas* are commonly associated with cyanobacteria (Morris et al. 2008, Biller et al. 2015, Hou et al. 2018, Kearney et al. 2021) and algae (Shibl et al. 2020, Cao et al. 2021). *Alteromonas* strains are easily isolated and cultured, partly attributed to their rapid response to the availability of organic matter (McCarren et al. 2010, Tinta et al. 2023). Indeed, a single *Alteromonas* strain has been shown to be capable of metabolizing almost the entire labile pool of marine organic carbon (Pedler et al. 2014). Phylogenetic, genomic and evolutionary studies have highlighted how genetic traits are exchanged between *Alteromonas* strains through genomic islands and plasmids (e.g. (Ivars-Martinez et al. 2008, Lopez-Perez et al. 2012, Fadeev et al. 2016, López-Pérez et al. 2016, López-Pérez et al. 2017, Koch et al. 2020)). Some *Alteromonas* strains may also have biotechnological applications (e.g. (Mehta et al. 2014, Concórdio-Reis et al. 2021)). Therefore, *A. macleodii* constitutes a relevant model organism in marine microbiology and biological oceanography (Wietz et al. 2022). The type strain, ATCC 27126, was isolated from surface seawater near Hawaii, and its physiology has been characterized in some detail (Baumann et al. 1972, Baumann et al. 1984).

Here, we describe conceptual and technical aspects of the community curation effort, performed at the University of Haifa (Israel) in February 2023 (Figure 1A). We specifically discuss the type of evidence typically available for emerging model organisms (Figure 1B), and then describe the curation of phenotypic traits with relevance for the ecological dynamics of *A. macleodii* (Figure 1C). We focused on pathways related to the uptake and utilization of carbon and nitrogen sources: 1) polysaccharides and one carbon (C1) compounds, as well as mixed acid fermentation; 2) nitrate, nucleotides and amino acids.

**Figure 1:**
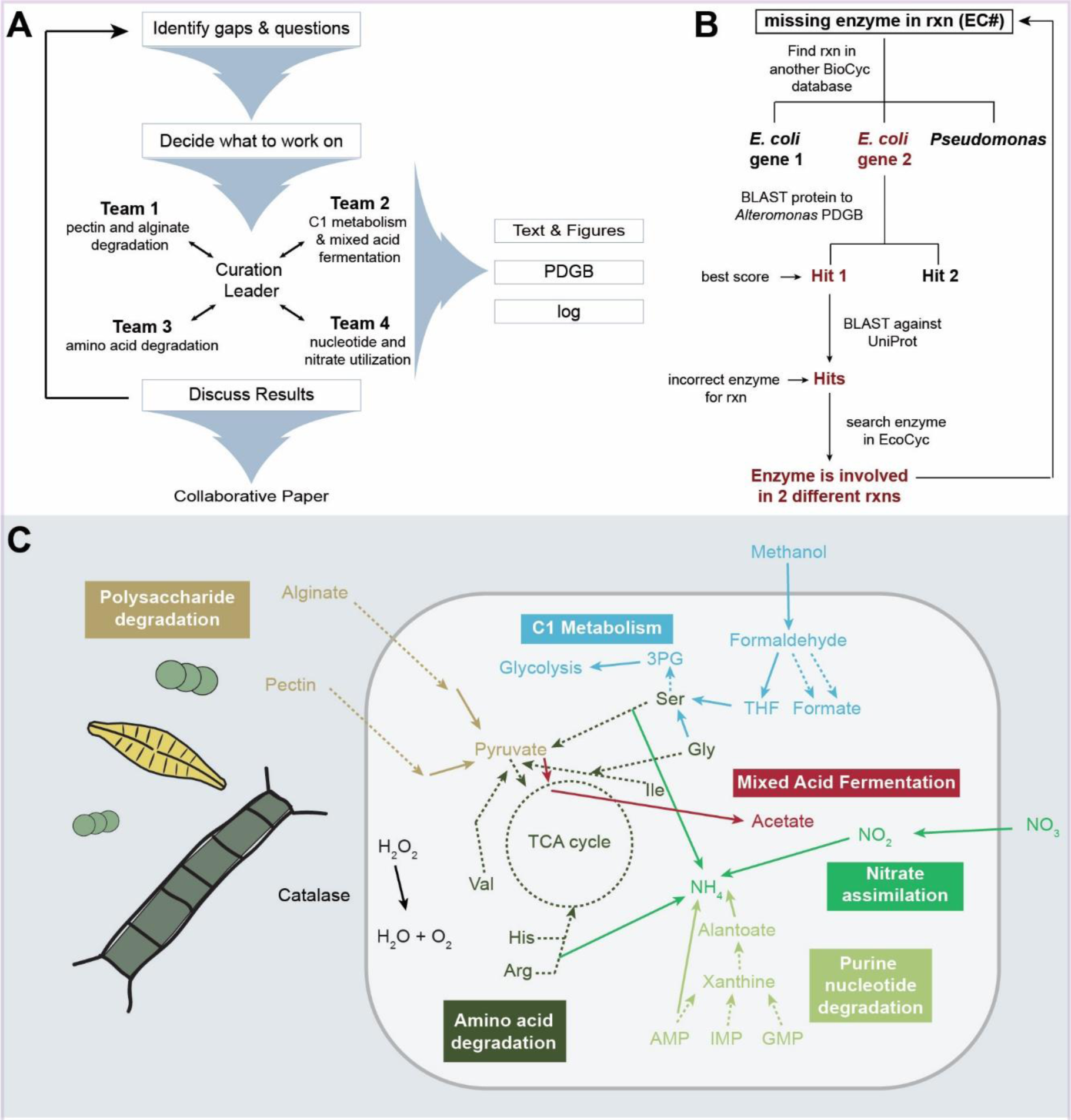
Schematic illustration of the A) workshop structure, B) key curation methodology, and C) main pathways curated.

### Metabolic curation − conceptual and technical aspects (Materials and Methods)

#### Automated reconstruction by PathwayTools

The starting point for the collaborative, manual curation process was a “Tier 3” PGDB, computationally derived by PathwayTools (Karp et al. 2021). Briefly, PathwayTools builds upon a genome annotated with the NCBI RefSeq PGAP pipeline (Li et al. 2020) for consistency between PGDBs, but RAST (Aziz et al. 2008), PROKKA (Seemann 2014) or DOE’s JGI/IMG (Markowitz et al. 2009) have also been used. The PathoLogic component of PathwayTools then generates the reactome, i.e. the set of enzyme-linked reactions, by mapping genes to enzymatic reactions in the MetaCyc reference knowledgebase (Caspi et al. 2020) based on a combination of gene and/or gene product names, Enzyme Commission (EC) number and Gene Ontology (GO) terms, if included in the annotation (Karp et al. 2021). This evidence then allows inferring the presence of specific metabolic pathways, based on a likelihood score that considers the fraction of enzymes identified per pathway, the presence of pathway-specific enzymes, and the expected phylogenetic distribution (e.g. a plant-specific pathway suggested to be in a bacterium would be flagged and the score penalized). Only pathways above a defined threshold are considered. As genes with unknown functions are common in model organisms (e.g. 35% in *E. coli*; (Ghatak et al. 2019)) and even more in non-model organisms (up to 80%; (Zoccarato et al. 2022)), the pathway thresholding step is intentionally permissive. This allows pathways to be integrated into the predicted metabolic network even if enzymes are missing. Next, a pathway hole filler (PHFiller) within PathoLogic identified missing reactions, and attempts to fill these using a BLAST search with multiple candidate genes from UniProtKB (Green et al. 2004). The resulting PDGB includes a report showing the score and completeness for each pathway, pathway holes that were “filled”, and pathways with remaining holes. This enables assessing the quality of the metabolic reconstruction by the curators, and advises where to perform manual curation − representing a robust quality control of the predicted metabolic network, where available experimental evidence is added via comments and evidence codes.

#### Community curation

Manual curation of the ATCC 27126 metabolic network was mostly performed during a four-day workshop by diverse researchers, including graduate students, postdocs and PIs under the guidance of a BioCyc curator (Lisa Moore) (Figure 1A). All workshop participants are co-authors on this paper. Prior to the curation workshop, a five-day course introduced the fundamentals of metabolic reconstruction and downstream uses, e.g. interpreting ‘omics data in light of metabolism. The workshop participants decided on the priorities for curation, taking into account the research interest in *Alteromonas* as versatile utilizers of dissolved and particulate organic matter, and their interactions with phytoplankton. Subteams of 3-5 curators focused on the curation of one or more pathways using the BioCyc web tool (unpublished). Often, multiple pathways were combined for visual interpretation using the “Pathway collage” tool in BioCyc (Paley et al. 2016).

#### Evidence types

Most organisms whose PDGBs undergo manual curation are widely studied; often being genetically tractable and/or medically-important taxa. Such organisms usually have accompanying gene-specific information, such as knock-out phenotypes or biochemical assays with purified proteins. In contrast, emerging model organisms often lack such information, and may not be genetically tractable (e.g. *Prochlorococcus* strains MED4 and SS120 with Tier-2 PDGBs available in BioCyc). ATCC 27126 was initially described in 1972 (Baumann et al. 1972), yet knock-out phenotypes have been only described for genes encoding a nitrate reductase and siderophore synthesis proteins (Diner et al. 2016, Manck et al. 2022). As a result, we considered additional types of evidence for metabolic reconstruction. Firstly, we compiled a list of media on which ATCC 27126 can grow, based on published studies as well as experiments performed for the curation workshop (Supplementary Excel File). Since ATCC 27126 can grow on minimal media with C, N, P and Fe sources but without amino acids, vitamins or cofactors, complete pathways for producing these compounds must be present. Such information was added as metadata to the specific pathway descriptions in the PGDB. Secondly, we considered evidence from related *Alteromonas* strains. For example, polysaccharide utilization pathways were curated through comparison with *A. macleodii* 83-1, a model polysaccharide degrader with 98% average nucleotide identity to ATCC 27126 (Koch et al. 2019). Finally, we identified candidate genes filling a specific pathway hole using Reciprocal Best BLAST (RBBH, (Altenhoff et al. 2009), Fig. 1B), using candidate “hole filling” genes identified in MetaCyc or EcoCyc using the EC number for each missing reaction. The protein sequence (from *E. coli* or, if using MetaCyc, from the closest relative of *Alteromonas*) was then queried against the PGDB using BLASTP within BioCyc. The best hit in ATCC 27126 was queried against the UniProtKB/Swiss-Prot database (The_UniProt_Consortium 2022). A gene product in ATCC 27126 was considered as RBBH (i.e. fill a pathway hole) if the best hit in UniProtKB/Swiss-Prot was annotated as the same function or EC number as the initial MetaCyc/EcoCyc query. In some cases, additional information was considered, such as the specificity of the annotation (e.g. methanol dehydrogenase vs. dehydrogenase), or BLAST sequence similarity and query cover. For Figure 3C, multiple sequence alignments were performed using MAFFT (Katoh et al. 2013). Phylogenetic analyses were also utilized, and maximum likelihood trees of *Alteromonas* alcohol dehydrogenases and 16S rRNA genes aligned with MUSCLE in AliView were inferred using IQ-TREE v1.5.4 (Edgar 2004, Larsson 2014, Nguyen et al. 2014).

## Results and discussion

The metabolic curation of *A. macleodii* ATCC 27126 was performed using a Tier-3 PDGB generated from NCBI genome assembly GCF_000172635.2. The reactions and pathways were then curated as described above, resulting in a Tier-2 PDGB (Table 1). The supplementary Excel File provides a detailed log of all steps within the collaborative curation. Below we discuss each of the main pathways or processes curated (Figure 1C).

**Table 1.** Summary statistics of the curated Tier-2 PGDB of *Alteromonas macleodii* ATCC 27126.

|  |  |
| --- | --- |
| Genes: | 3,962 |
| Pathways: | 244 |
| Enzymatic Reactions: | 1,492 |
| Transport Reactions: | 19 |
| Polypeptides: | 3,829 |
| Protein Complexes: | 49 |
| Enzymes: | 915 |
| Transporters: | 304 |
| Compounds: | 1,029 |
| Transcription Units: | 2,659 |
| tRNAs: | 52 |
| Protein Features: | 6,181 |
| GO Terms: | 4 |

## Carbon sources

### Carbohydrate-active enzymes and polysaccharide degradation

ATCC 27126 encodes several pathways to degrade algal polysaccharides, important bacterial nutrient sources in the oceans. Degradation relies on polysaccharide lyases (PL), glycoside hydrolases (GH) and carbohydrate esterases (CE). These genes are mostly encoded in polysaccharide utilization loci (PULs), operon-like gene clusters with concerted regulation. Here, using complementary evidence, we curated the pathways for pectin and alginate degradation (Figure 2). We annotated genes encoding carbohydrate-active enzymes (CAZymes, (Zhang et al. 2018, Drula et al. 2022)) in ATCC 27126 by comparison with transcriptomic and proteomic data from a closely related strain, *A. macleodii* 83-1. Both strains harbor homologous alginolytic and pectinolytic PULs (Neumann et al. 2015, Koch et al. 2019), which are significantly upregulated in 83-1 when growing with an alginate and pectin mix (Koch et al. 2019).

**Figure 2:**
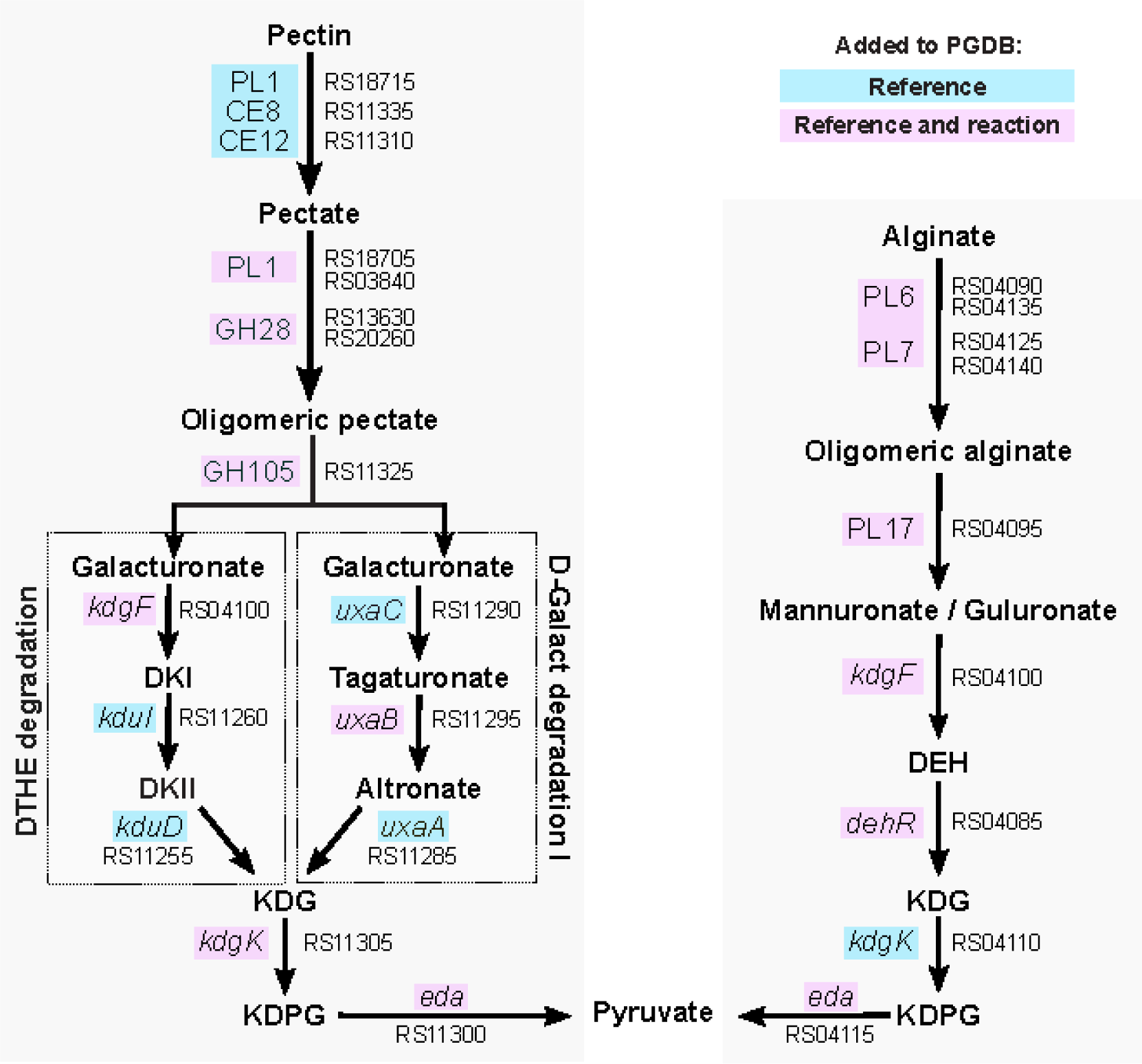
Curated pectin and alginate degradation pathways. The pectin superpathway (left; PWY2OKO-5) encompasses initial depolymerization and demethylation followed by 4-deoxy-L- threo-hex-4-enopyranuronate degradation (PWY-6507; abbreviated DTHE) or D-galacturonate degradation I (GALACTUROCAT-PWY; abbreviated D-Galact) for unsaturated and saturated galacturonates respectively. Both pectin and alginate degradation (right; PWY-6986-1) eventually result in KDG, KDGP and pyruvate, but these metabolites are generated via dedicated enzymes. DKI: 5-keto-4-deoxyuronate; DKII: 2,5-diketo-3-deoxygluconate; DEH: 4-deoxy-l-erythro-5- hexoseulose uronate; KDG: 2-keto-3-deoxygluconate; KDPG: 2-dehydro-3-deoxy-D-gluconate 6- phosphate; us: unsaturated. Blue boxes indicate genes for which the reference information was updated in the PDGB. Pink boxes indicate genes for which new reactions also were added. Gene names and locus tags are shown for each reaction.

For pectin degradation, we imported relevant degradation subpathways from MetaCyc and constructed a new pectin superpathway, PWY2OKO-5, encompassing depolymerization (via PL1, GH28 and GH105) and demethylation (via CE8 and CE12) (Figure 2). The resulting galacturonates are then processed via 4-deoxy-L-threo-hex-4-enopyranuronate (PWY-6507) and D-galacturonate (GALACTUROCAT-PWY) pathways respectively, which we added to the ATCC 27126 PDGB. The released methanol is possibly metabolized by alcohol dehydrogenases (see below).

Curating the alginate degradation pathway benefited from RT-qPCR evidence in ATCC 27126, showing significantly higher expression of PL6 and PL7 lyases with alginate as sole nutrient source (Neumann et al. 2015). Biochemical assays in 83-1 with cloned, homologous enzymes confirmed alginate lyase activity, and characterized salinity and temperature optima (Gerlach 2017)). However, structural elucidation failed, since not enough soluble enzyme was obtained (Gerlach 2017). Our curation process involved adding reactions for 4-deoxy-l-erythro-5- hexoseulose uronate (DEH) reductase (alginate degradation) as well as *kdgF* (both pathways), since uronate conversion does not occur spontaneously (Hobbs et al. 2016) as originally annotated in BioCyc (Figure 2).

Both pectin and alginate are composed of uronate sugars, eventually yielding pyruvate from 2- keto-3-deoxygluconate (KDG) and 2-dehydro-3-deoxy-D-gluconate 6-phosphate (KDPG, Figure 2). Notably, KDG and KDPG are generated via pectin- or alginate-specific *kdgK* and *eda* genes encoded in the respective PULs for each polysaccharide. ATCC 27126 also encodes another *eda* copy (MASE_RS11155) not induced by pectin or alginate in 83-1, which might be a “generic” variant to convert KDPG from other sources.

### C1 metabolism

One-carbon (C1) and methylated compounds are important bacterial substrates in the marine environment (Dixon et al. 2013, Lidbury et al. 2014). Here we examined the potential of ATCC 27126 to metabolize methanol, formaldehyde (the central C1 intermediate), formate, and related cofactors and enzymes (Figure 3A).

**Figure 3:**
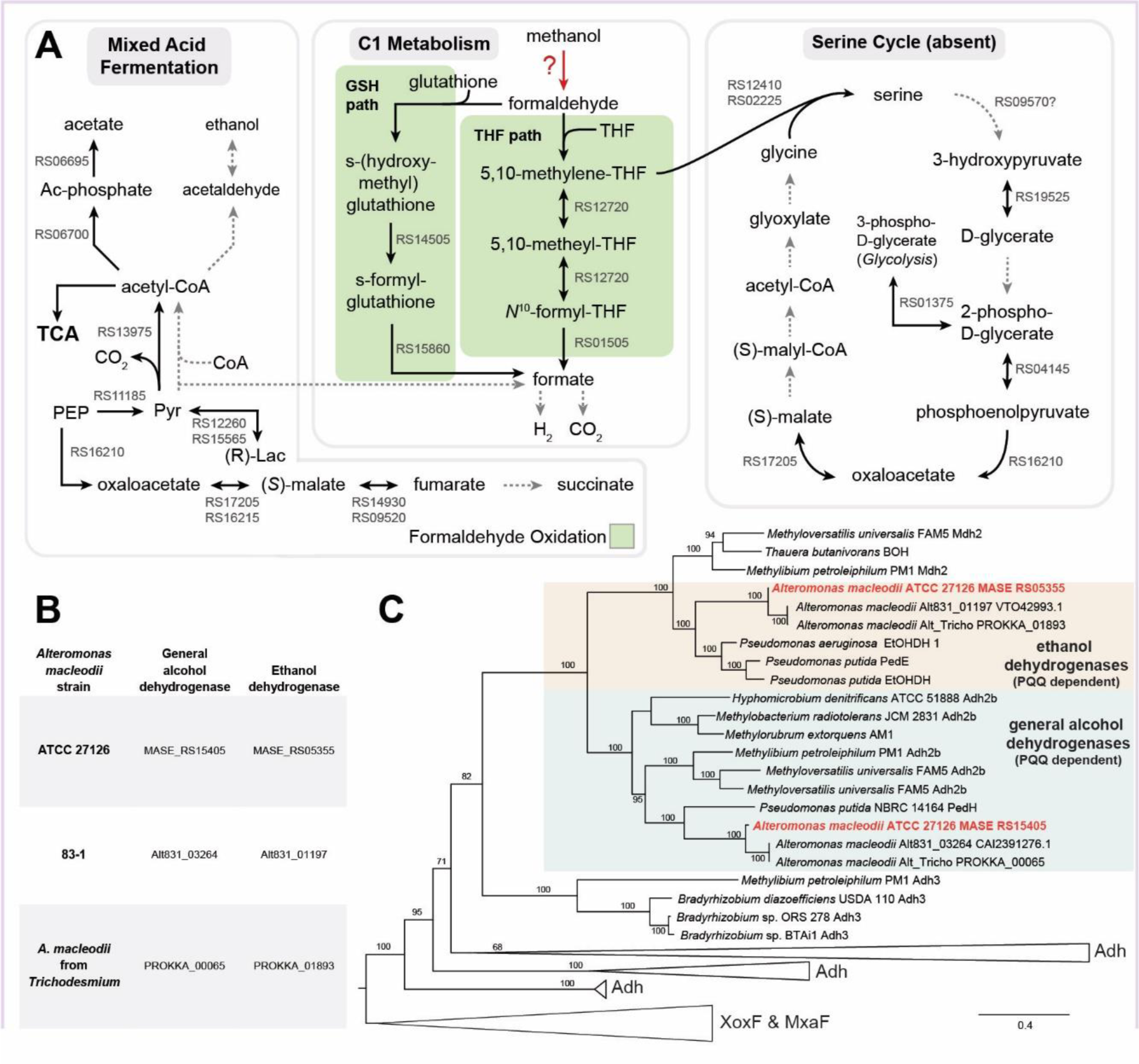
C1 metabolism, the serine cycle and mixed acid fermentation in *A. macleodii* ATCC 27126. A) Potential routes for one-carbon and methylated compounds. Bold arrows represent reactions identified in the ATCC 27126 genome, dotted arrows are missing reactions. THF: Tetrahydrofolate; Lac: lactate; Pyr: pyruvate; PEP: phosphoenolpyruvate. B) Putative methanol/ethanol dehydrogenase genes in *Alteromonas* strains that grow and do not grow on methanol. C) Maximum likelihood tree (IQ-TREE) inferred under the Q.pfam+F+I+G4 model from alcohol dehydrogenase proteins from ATCC 27126 and other strains. Support values represent 100 bootstrap pseudoreplicates.

#### Methanol/Ethanol Dehydrogenases

Methanol is commonly produced by phytoplankton and cyanobacteria (Mincer et al. 2016), but only some *A. macleodii* strains can grow on methanol. Several *A. macleodii* strains were isolated from *Trichodesmium* using media with methanol as the sole carbon source, attributed to pyrroloquinoline quionone (PQQ)-dependent alcohol dehydrogenases (ADHs) encoded in their genomes (Lee et al. 2017). In contrast, neither ATCC 27126 nor 83-1 grow on methanol (Baumann et al. 1984, Koch et al. 2019), although both encode the same PQQ-dependent ADH genes (Figure 3B). ATCC 27126 and 83-1 also encode an operon (*pqqABCDE*) with the dehydrogenase PQQ cofactor, which was identified using mass spectrometry in 83-1 (Koch et al. 2019). Our phylogenetic analysis of predicted PQQ-dependent ADH genes did not support their potential role in methanol oxidation (Figure 3C): MASE_RS15405 is within a clade of general ADHs that could potentially mediate methanol oxidation, while MASE_RS05355 is more related to ethanol dehydrogenases. Additionally, both genes show only moderate amino acid identity (30%) with the known PQQ-dependent methanol dehydrogenases XoxF and MxaF (Keltjens et al. 2014). These ADHs might alternatively convert ethanol to acetaldehyde; indeed, ATCC 27176 was shown to grow on ethanol (Baumann et al. 1984).

ATCC 27126 also encodes a zinc-dependent ADH (MASE_RS02430) and three iron-containing ADHs (MASE_RS06555, MASE_RS01350, MASE_RS11390). Iron- and zinc-containing ADHs can catalyze the oxidation of methanol, ethanol, and other alcohols (e.g. (Vries et al. 1992, Antoine et al. 1999, Liu et al. 2009)), although some of these enzymes may preferentially catalyze the reverse reaction (i.e. reduction; see mixed acid fermentation below). Alternatively, both ethanol and methanol can be converted to acetaldehyde and formaldehyde, respectively, during hydrogen peroxide detoxification by catalase and related enzymes (e.g. (Ro et al. 2003)). ATCC 27126 encodes five catalase genes that potentially mediate interactions with phytoplankton (Hennon et al. 2017), yet it is unclear whether this pathway (primarily for detoxification) produces significant energy to support growth.

In summary, despite finding several candidate genes for methanol oxidation, it remains unclear why ATCC 27126 does not grow on methanol while other strains do. Potentially, ATCC 27126 can metabolize methanol, but not grow on it as a sole C source. The same has been shown for *Pelagibacter,* which encodes multiple genes involved in C1 metabolism (including ADHs), oxidizes methanol and other C1 compounds to CO2, yet does not incorporate C1 compounds into biomass (Sun et al. 2011). Therefore, varying abilities between *A. macleodii* strains in their ability to utilize methanol may be due to different downstream oxidation steps.

#### Formaldehyde Metabolism

Formaldehyde is a common byproduct of methanol oxidation and a critical intermediate in C1 metabolism. It is also often cytotoxic and must be metabolized quickly. ATCC 27126 is predicted to encode two parallel routes for formaldehyde oxidation: the tetrahydrofolate-based (THF) and the glutathione-based (GSH) pathway (Figure 3A).

The THF pathway comprises four steps leading to formate; the first step is spontaneous (Kallen et al. 1966, Marx et al. 2003a) and the remaining three catalyzed (Müller et al. 2015). The product of the first step, 5,10-methylenetetrahydrofolate, can either enter the serine cycle (as in the methylotroph *Methylorubrum extorquens*, (Marx et al. 2003b)) or be converted into formyltetrahydrofolate (as in the facultative methylotroph *Bacillus methanolicus,* which lacks the serine cycle (Müller et al. 2015)). As discussed below, ATCC 27126 likely does not encode the serine cycle, suggesting the presence of dissimilatory formaldehyde conversion via the THF pathway, yielding formate.

The GSH pathway involves NAD, glutathione-dependent formaldehyde dehydrogenase (GSH- FDH, also called S-(hydroxymethyl)glutathione dehydrogenase), and S-formylglutathione hydrolase (FGH). S-(hydroxymethyl)glutathione, formed spontaneously by formaldehyde and glutathione, is the preferred *in vitro* and presumed *in vivo* substrate for GSH-FDH (Barber et al. 1996, Harms et al. 1996). Both GSH-FDH and FGH are encoded in the ATCC 27126 genome (MASE_RS14505 and MASE_RS15860, respectively). ATCC 27126 may need both THF and GSH pathways for C1 metabolism and/or detoxification of formaldehyde, as in *Methylorubrum extorquens* (Marx et al. 2003b).

#### Formate metabolism

Formate from formaldehyde oxidation is usually oxidized to CO2 (Maia et al. 2016), catalyzed by a formate hydrogen lyase complex (FHL) comprised of formate dehydrogenase *fdhF* and six subunits of hydrogenase 3. A BLAST search of *E. coli fdhF* (UniProt:P07658) showed several homologs in marine bacteria (mainly *Shewanella*), but no clear homolog in *Alteromonas*. An alternative route is the formate dehydrogenase operon (FDH), but we found only one of the four FDH genes (*fdhD*) in the ATCC 27126 genome. Although *fdhD* is encoded adjacent to another gene (MASE_RS07820) distantly related to a formate dehydrogenase (∼31% identical to *fdhH* from *E. coli*), MASE_RS07820 is a pseudogene due to a frame shift. Therefore, ATCC 27126 presumably cannot oxidize formate to CO2.

Formate also participates in the glutamylation of tetrahydrofolate. Glutamylated folate cofactors are required in various C1 reactions, acting as carriers of one-carbon units (Shane 1989). However, the first enzyme in this reaction, formate-tetrahydrofolate ligase, is not encoded in the ATCC 27126 genome. The lack of key metabolic pathways supports the observation that *A. macleodii* cannot grow with formate as a sole carbon source (Baumann et al. 1972). Future work could determine whether ATCC 27126 excretes formate, similar to some methylotrophs (Baev et al. 1992) and cyanobacteria (Heyer et al. 1991, Bertilsson et al. 2005, Sosa et al. 2019), potentially providing a carbon source for co-occurring organisms.

#### Serine Cycle

Formaldehyde can also be assimilated via the serine cycle (formaldehyde assimilation I pathway in BioCyc), yielding several intermediates for central carbon metabolism ((Anthony 2011), Figure 3A). In the first step, hydroxymethyltransferase (GlyA) catalyzes the reaction of 5,10-methylenetetrahydrofolate with glycine to form serine. ATCC 27126 encodes two GlyA proteins (MASE_RS12410 and MASE_RS02225) along with a putative serine-glyoxylate transaminase (MASE_RS09570), an enzyme that mediates the following conversion of serine to 3-hydroxypyruvate. Hydroxypyruvate reductase to reduce 3-hydroxypyruvate to D-glycerate, was not identified in ATCC 27126. Instead, we found 2-hydroxyacid dehydrogenase (MASE_RS19525), which also converts 3-hydroxypyruvate to D-glycerate. Genes encoding the remaining essential enzymes were not detected, including EC 6.2.1.9 and EC 4.1.3.24 that define the presence of the serine cycle in methylotrophs. This genomic evidence, along with the lack of growth on methanol as a sole carbon source (Koch et al. 2019) supports the absence of the serine cycle in ATCC 27126.

## Mixed acid fermentation

Mixed acid fermentation involves the catabolism of pyruvate to lactate, formate, acetate, ethanol and succinate when no exogenous electron acceptors are available. Marine particles, to which *Alteromonas* is often attached (e.g. (Mestre et al. 2017, Roth Rosenberg et al. 2021), potentially have microaerobic or anaerobic micro-niches (Bianchi et al. 2018), yet. ATCC 27126 has been described as strictly aerobic (Baumann et al. 1972). Mixed acid fermentation was inferred in ATCC 27126 during the computational reconstruction of PDGB, albeit with pathway holes. Hence, we decided to investigate this pathway further.

ATCC 27126 encodes 7 out of 11 enzymes for mixed acid fermentation, including those catalyzing the conversions of acetyl CoA to acetyl phosphate and acetate, pyruvate to lactate, as well as the formation of fumarate from phosphoenolpyruvate (PEP) (Figure 3A). The presence of these reactions is supported by detecting acetate, lactate and succinate in extracellular polysaccharides (EPS), where they act as non-carbohydrate substituents (Raguénès et al. 2003, Concórdio-Reis et al. 2021). However, ATCC 27126 does not encode a fumarate reductase enzyme. Thus, although traces of succinyl have been reported in *Alteromonas* EPS (Concórdio-Reis et al. 2021) and fumarate reduction can principally occur via succinate dehydrogenase (Cecchini et al. 2002), presence of mixed-acid fermentation in ATCC 27126 remains unclear. Moreover, ATCC 27126 lacks the genes catalyzing the initial anaerobic conversion of pyruvate and CoA to acetyl-CoA and formate (*pflB* and *tdcE* in *E. coli* K-12). As discussed above, it also lacks the formate dehydrogenase complex that catalyzes the sequential conversion of formate to CO2. Finally, ATCC 27126 lacks the canonical genes catalyzing the reduction of acetyl-CoA to acetaldehyde and ethanol. While, in principle, acetaldehyde could be reduced to ethanol by one of the PQQ- or zinc-dependent alcohol dehydrogenases (described above) working in reverse, the lack of an acetaldehyde dehydrogenase gene suggests that this part of mixed acid fermentation may be dysfunctional. Therefore, the bioinformatic evidence for the full mixed acid fermentation process is inconclusive. Nonetheless, many of these reactions also work in reverse (e.g., lactate dehydrogenase), and may enable ATCC 27126 to catabolize organic acids excreted by co-occurring algae even under oxic conditions (Bertilsson et al. 2005, Braakman et al. 2017). Indeed, ATCC 27126 can grow on lactate or pyruvate as sole carbon sources ((Baumann et al. 1984), Supplementary Excel file.

## Nitrogen sources

### Amino acid degradation

Amino acids, constituting a significant fraction of organic nitrogen in the oceans (Lee et al. 2000), can serve as both nitrogen and carbon sources. ATCC 27126 grows well with peptides or a mixture of amino acids as sole carbon sources (Forchielli et al. 2022). Although comparative genomics suggested that most *Alteromonas* spp. can degrade almost every amino acid (Figure 4A), ATCC 27126 growth was only reported on alanine and glycine (Baumann et al. 1984). We asked whether this inconsistency is due to missing genes in the predicted amino acid degradation pathways as they enter the TCA cycle (Figure 4B). For seven pathways with putative holes, we identified candidate genes for three pathways that could “close” these holes using RBBH (Supplementary Excel File). The putative “hole-filling” genes MASE_RS07620 and MASE_RS07610 are encoded within a predicted operon for L-leucine degradation, somewhat similar to the *liu* operon from *Pseudomonas aeruginosa* (Figure 4C, (Kazakov et al. 2009)). The predicted “hole-filling” gene MASE_RS01650 is part of an operon for arginine degradation. However, we could not reconstruct the pathways for the degradation of threonine, tryptophan and tyrosine. Notably, there are holes in the pathways for tryptophan and tyrosine degradation in *E. coli* SIJ488, yet this strain can utilize these amino acids as sole nitrogen sources (Schulz-Mirbach et al. 2022). Therefore, these pathways might still be functional in ATCC 27126, although further experimental work is needed to test this hypothesis.

**Figure 4.**
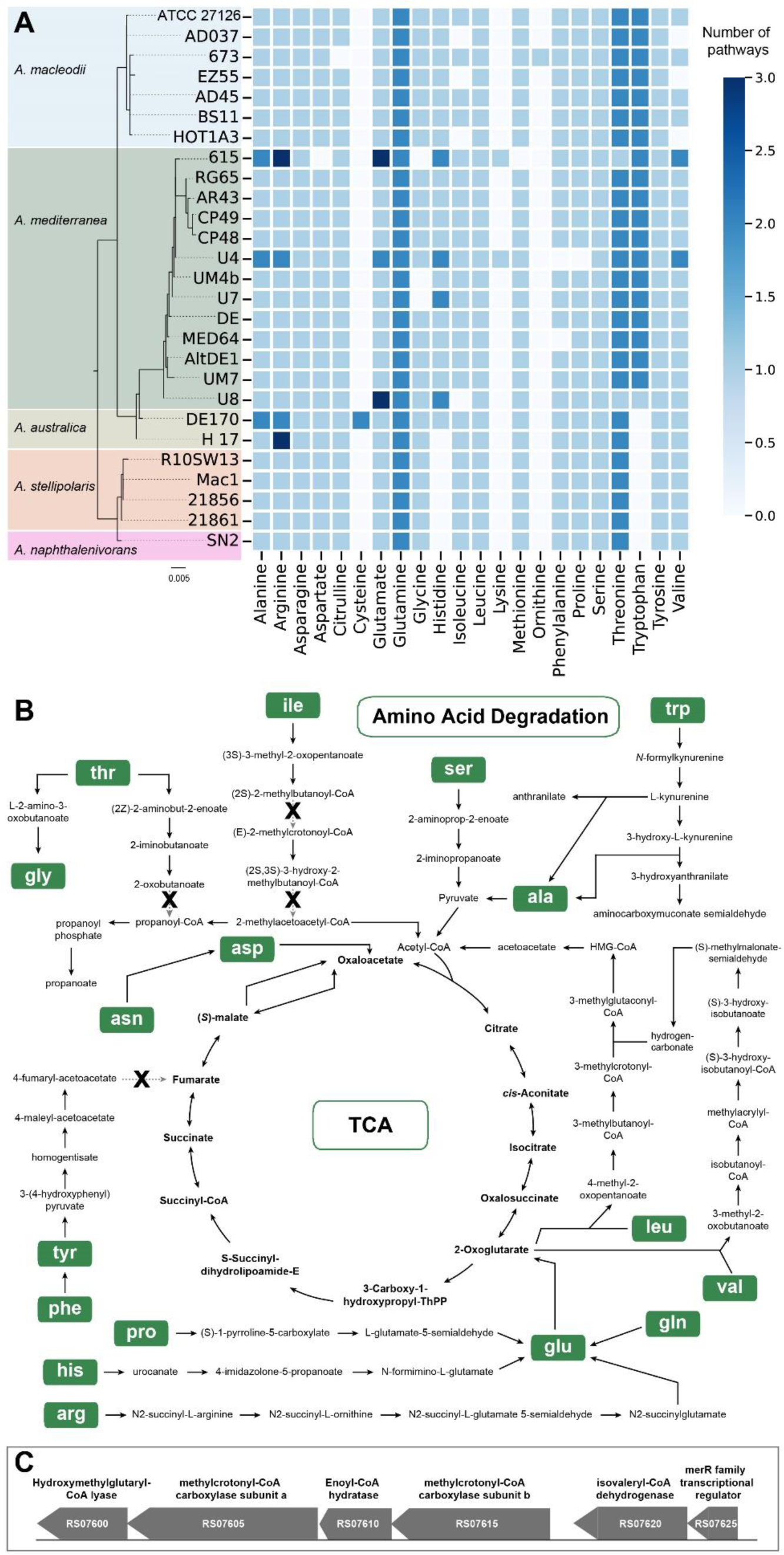
Amino acid degradation pathways and their holes. A) Number of predicted amino acid degradation pathways per genome across *Alteromonas* spp., determined using BioCyc’s Comparative Genomics tool. The taxa are ordered based on a maximum likelihood tree (IQTREE) inferred under the TIM3+F+I model from full-length 16S rDNA. B) Amino acid utilization pathways in ATCC 27126, drawn based on an overview from the BioCyc “Pathway Collage” tool. Missing reactions (“pathway holes”) are highlighted by an X. Methionine degradation is not shown as it enters the TCA cycle via multiple other pathways. C) Predicted operon for branched chain amino acid degradation, with similarities to the *liu* operon in *Pseudomonas aeruginosa* (Kazakov et al. 2009).

### Nucleotide degradation

Purine nucleotides can serve as nitrogen sources for bacteria (Huang et al. 2022). The purine nucleotide degradation II superpathway in MetaCyc is composed of three sequential pathways: 1) purine nucleotide degradation II (starting with AMP, GMP and IMP each yielding urate); 2) urate conversion to allantoin; and 3) allantoin degradation (Figure 5A). The first pathway is fully present in ATCC 27126 (Figure 5A), whereas the second was not predicted despite finding genes encoding 2 out of 3 reactions. However, RBBH using the *puuD* gene from *Agrobacterium fabrum* identified MASE_RS07125 as the putative hole-filling gene in the urate conversion to allantoin pathway, encoding a protein containing a urate oxidase domain (PF016181) (Figure 5B). We therefore curated the *puuD* gene, and added the missing urate conversion pathway.

**Figure 5:**
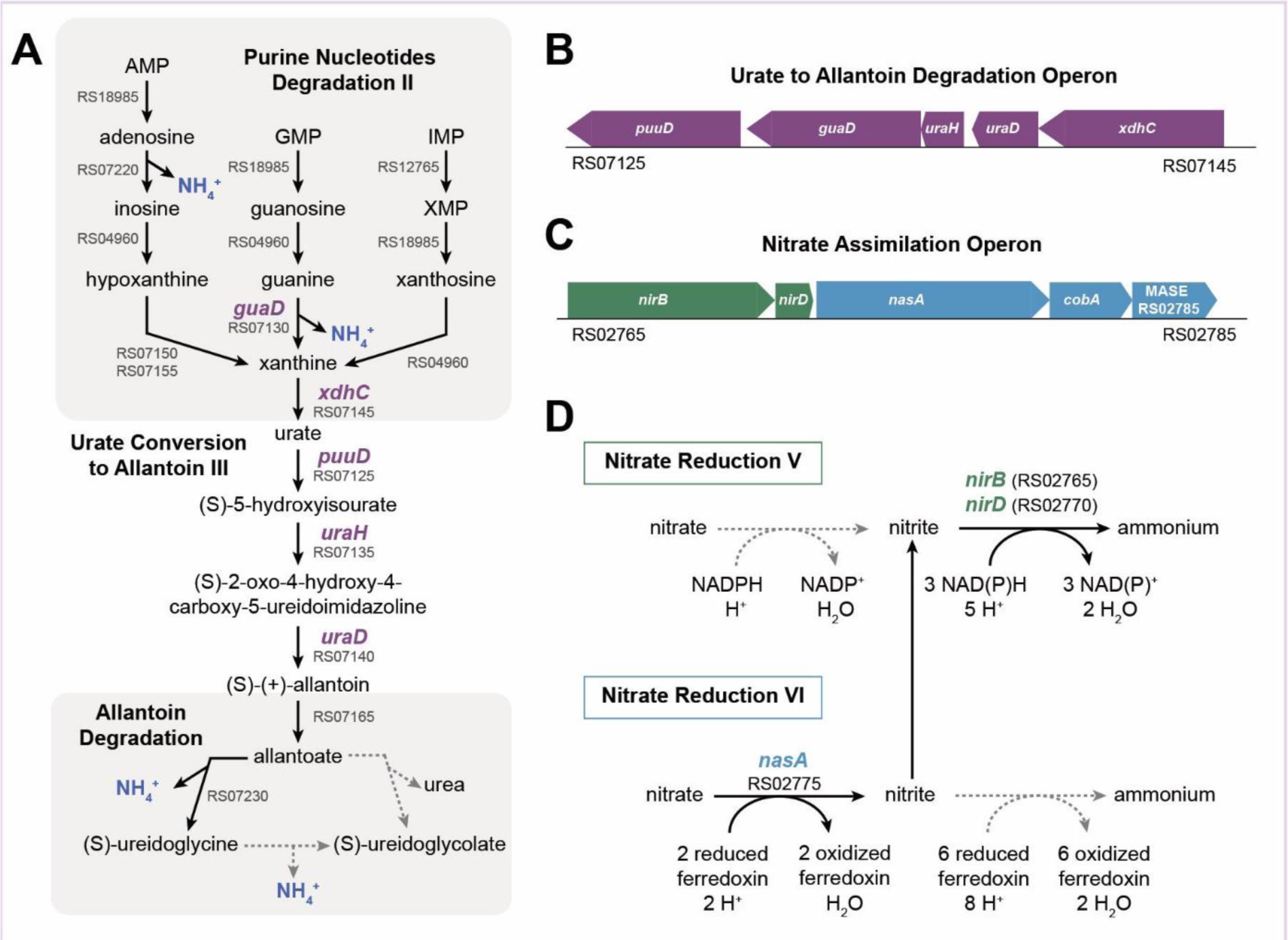
Nucleotide and nitrate utilization pathways. A) The purine nucleotide degradation II (aerobic) superpathway in ATCC 27126. Dotted arrows show missing reactions, stages where NH4 is released are highlighted. B) Genomic region surrounding the putative urate to allantoin degradation operon (*puuD* homologue). C) Putative assimilatory nitrate reductase operon. D) Suggested hybrid nitrate reductase pathway in ATCC 27126, utilizing different cofactors for nitrate and nitrite reduction.

Further metabolism of allantoin to glyoxylate can occur via S-ureidoglycine or S-ureidoglycolate (Figure 5A). However, these pathways are incomplete in ATCC 27126, and we were unable to identify hole-filling genes. Accordingly, ATCC 27126 cannot grow on allantoin as sole C source (Baumann et al. 1984). Taken together, the missing allantoin degradation pathway might explain why ATCC 27126 cannot grow on nucleotides as sole carbon source (Baumann et al. 1984). Nevertheless, there is a possibility that this strain can use purines as N sources, since ammonium is released at multiple steps of the degradation pathway. If so, we predict ATCC 27126 to excrete allantoin, allantoate and/or S-ureidoglycolate during nucleotide degradation.

### Nitrate assimilation

ATCC 27126 can grow on nitrate as a sole N source, likely incorporated to biomass via assimilatory reduction (Diner et al. 2016). The draft PDBG did not include assimilatory, but rather dissimilatory, nitrate reduction (i.e. denitrification), via a cluster of one nitrate reductase (*nasA*; MASE_RS02775) and two nitrite reductase genes (*nirBD*, MASE_RS02765 and MASE_RS02770; Figure 5C). A *nasA* mutant of ATCC 27126 cannot grow on nitrate, highlighting that this operon encodes assimilatory nitrate reduction (Diner et al. 2016). Additionally, there is no evidence for denitrification (Baumann et al. 1984), although distantly related Alteromonads may respire nitrate (Moisander et al. 2018). Therefore, dissimilatory nitrate reduction was removed from the PDGB.

The putative assimilatory nitrate reduction pathway in ATCC 27126 shows an unusual combination of genes and required cofactors (Figure 5D). In most bacteria and fungi, nitrate and nitrite reductases use NAD(P)H as electron donor (Lin et al. 1997), whereas cyanobacteria use ferredoxin (Herrero et al. 1997). The ATCC 27126 *nasA* is most similar to homologs of the nitrate reduction V pathway common to many bacteria and fungi (BLAST bit score 678). However no homolog was found for the *nasC* protein, often associated with *nasA* to form the nitrate reduction complex (Lin et al. 1997). The second best hit for *nasA* was to a cyanobacterial nitrate reductase *narB* (EC 1.7.7.2, BLAST bit score 607 with *narB* from *Synechococcus elogantus*), which does not require additional subunits, is closer in length and shares a ferredoxin-binding domain and 3 molybdopterin-containing Pfam domains with *A. macleodii* ATCC 27126 *nasA*. We therefore propose that ATCC 27126 encodes a “chimeric” nitrate assimilation pathway, with a ferredoxin-utilizing homolog of the cyanobacterial nitrate reductase *narB* followed by NAD(P)H-utilizing bacterial homologs of the nitrite reductase *nirBD* complex. This hypothesis requires experimental verification.

### Conclusions and future prospects

Our interactive community curation contributes to the metabolic reconstruction of *Alteromonas macleodii* ATCC 27126 by combining bioinformatic, genomic and physiological evidence, together with an extensive literature review. This approach explored specific traits related to *Alteromonas* metabolism and ecology and provides recommendations for future experimental work. Formalizing this curation as a Tier-2 PDGB in BioCyc (PDGB ID: 2OKO, Nov 2023 version available freely on DOI https://github.com/Sher-lab/amac/) helps consolidating the biological understanding of *A. macleodii*, providing a dynamic and growing “metabolic encyclopedia”.

ATCC 27126 and other *A. macleodii* strains can utilize complex polysaccharides, but cannot grow on methanol or formate (Baumann et al. 1972, Baumann et al. 1984). The fate of pectin-derived methanol remains unclear, since the specificity of encoded ADHs could not be determined. If methanol can be metabolized to formaldehyde, the latter may either be fed into glycolysis for energy (but not biomass) production through a partial serine cycle or metabolized to formate which is probably excreted into the environment. Taken together, *A. macleodii* might hence show a “sharing” phenotype during pectin degradation, releasing simple organic compounds that can be metabolized by other bacteria (Fritts et al. 2021).

Some of the evidence supports the finding of *A. macleodii* in certain niches, but not in others. For example, polysaccharides degradation might support its association with algae and polymer microgels (Mitulla et al. 2016). Similarly, growth on peptides (Forchielli et al. 2022), together with their ability to utilize various forms of dissolved organic phosphorus (Srivastava et al. 2021), may support their growth on other forms of organic matter such as dead jellyfish biomass (Tinta et al. 2023) or deep-sea particles (Zhao et al. 2020). In contrast, ATCC 27126 cannot grow on chitin, the most abundant polysaccharide in zooplankton (Baumann et al. 1984), and indeed *Alteromonas* are not part of the core copepod microbiome (Datta et al. 2018). Future curation efforts and accompanying experiments focusing on carbohydrate, protein and organophosphorus degradation may help clarifying the “metabolic niche” of *A. macleodii*, determine whether they are associated with specific organisms, and raise testable hypotheses regarding their spatial and temporal variability in the oceans (Alonso-Sáez et al. 2007, Roth Rosenberg et al. 2021).

The finding of *Alteromonas* on marine particles (Roth Rosenberg et al. 2021, Henríquez-Castillo et al. 2022, Wietz et al. 2022) might be connected to hypoxic or anoxic micro-niches (Bianchi et al. 2018), yet the evidence for fermentation in ATCC 27126 is inconclusive. We propose that a better characterization of the “oxygen niche” of *Alteromonas* will help to understand their involvement in particle colonization and degradation (Zhao et al. 2020). Similarly to understanding how *Alteromonas* utilizes amino acids, nucleotides or nitrate will contribute to understanding their role in nitrogen cycling, given that proteins can comprise 50-60% of phyto- and zooplankton biomass (Lee et al. 2000, Geider et al. 2002, Helland et al. 2003, Finkel et al. 2016, Givati et al. 2023). Importantly, comparing cultured strains with environmental datasets will need to consider the diversity within the *Alteromonas* clade (e.g. (Gonzaga et al. 2012, López-Pérez et al. 2016, López-Pérez et al. 2017, Koch et al. 2020).

Our community curation clarified important aspects of the physiology and ecology of ATCC 27126, and suggested relevant experimental directions. However, we also highlight key challenges in studying ecologically important but less described organisms. Only a few *A. macleodii* genes have been functionally verified, resulting in often indirect evidence for the presence or absence of specific reactions. Deciding whether or not to include pathways in the PDGB was therefore sometimes subjective, especially if requiring “hole filling”. Furthermore, manual curation requires in-depth knowledge of multiple aspects of metabolism, which is typically beyond the expertise of any single curator or a diverse group like in our study. We suggest that any curation process clearly records the evidence used to decide whether a pathway is present, enabling future users to revisit the metabolic reconstruction before generating genome-scale models. Secondly, there are no clear guidelines for the incorporation of genomic information for metabolic reconstructions. While the conservation of (partial) pathways in closely related organisms may suggest they are functional, our results for amino acid utilization highlight that such hypotheses are not always fully supported. Furthermore, systematically addressing the correlation between pathway phylogeny and function may facilitate metabolic curation by harvesting experimental results from yet-uncultured organisms, e.g. using metagenome-assembled or single-cell genomes (Tinta et al. 2023). Finally, a metabolic model widely accessible to the scientific community needs to be compatible with downstream analyses. There are currently several frameworks for representing cell metabolism, e.g. KEGG (Kanehisa et al. 2000) and modelSEED (Seaver et al. 2021), yet translating metabolic reconstructions from between frameworks can be difficult since terms and structure (e.g., where do they draw boundaries between pathways) are not compatible. Moreover, due to the cost of maintaining the BioCyc infrastructure, accessing most PDGBs requires payment. We hope that the insights from our community curation will provide incentives for research groups and funding agencies to include metabolic knowledge for environmentally important organisms in financially supported databases.

## Acknowledgements

This study was supported by grant 1786/20 from the Israel Science Foundation (to DS) and the University of Haifa Data Science Research Center, Rectors office and International School. AM, PK and LM were supported by funding from SRI. EEG was supported by the Simons Foundation Postdoctoral Fellowship in Marine Microbial Ecology (award ID: 993200). We thank Amanda Mackie for assistance with the annotation and Moran Altman for technical coordination.

## Author contributions

DS and LM designed the workshop and study; all authors performed bioinformatics analyses and curated the PDGB; DS, EG, MW, SG, LZ, OW and LM wrote the manuscript with input from all authors.

## References

Alonso-Sáez, L., V. Balagué, E. L. Sà, O. Sánchez, J. M. González, J. Pinhassi, R. Massana, J. Pernthaler, C. Pedrós-Alió and J. M. Gasol (2007). “Seasonality in bacterial diversity in north-west Mediterranean coastal waters: assessment through clone libraries, fingerprinting and FISH.” FEMS Microbiology Ecology 60(1): 98–112. 10.1111/j.1574-6941.2006.00276.x

Altenhoff, A. M. and C. Dessimoz (2009). “Phylogenetic and Functional Assessment of Orthologs Inference Projects and Methods.” PLoS Computational Biology 5(1): e1000262. 10.1371/journal.pcbi.1000262

Anthony, C. (2011). “How half a century of research was required to understand bacterial growth on C1 and C2 compounds; the story of the serine cycle and the ethylmalonyl-CoA pathway.” Sci Prog 94(Pt 2): 109–137. 10.3184/003685011×13044430633960

Antoine, E., J.-L. Rolland, J.-P. Raffin and J. Dietrich (1999). “Cloning and over-expression in Escherichia coli of the gene encoding NADPH group III alcohol dehydrogenase from Thermococcus hydrothermalis.” European Journal of Biochemistry 264(3): 880–889. 10.1046/j.1432-1327.1999.00685.x

Arkin, A. P., R. W. Cottingham, C. S. Henry, N. L. Harris, R. L. Stevens, S. Maslov, P. Dehal, D. Ware, F. Perez, S. Canon, M. W. Sneddon, M. L. Henderson, W. J. Riehl, D. Murphy-Olson, S. Y. Chan, R. T. Kamimura, S. Kumari, M. M. Drake, T. S. Brettin, E. M. Glass, D. Chivian, D. Gunter, D. J. Weston, B. H. Allen, J. Baumohl, A. A. Best, B. Bowen, S. E. Brenner, C. C. Bun, J.-M. Chandonia, J.-M. Chia, R. Colasanti, N. Conrad, J. J. Davis, B. H. Davison, M. DeJongh, S. Devoid, E. Dietrich, I. Dubchak, J. N. Edirisinghe, G. Fang, J. P. Faria, P. M. Frybarger, W. Gerlach, M. Gerstein, A. Greiner, J. Gurtowski, H. L. Haun, F. He, R. Jain, M. P. Joachimiak, K. P. Keegan, S. Kondo, V. Kumar, M. L. Land, F. Meyer, M. Mills, P. S. Novichkov, T. Oh, G. J. Olsen, R. Olson, B. Parrello, S. Pasternak, E. Pearson, S. S. Poon, G. A. Price, S. Ramakrishnan, P. Ranjan, P. C. Ronald, M. C. Schatz, S. M. D. Seaver, M. Shukla, R. A. Sutormin, M. H. Syed, J. Thomason, N. L. Tintle, D. Wang, F. Xia, H. Yoo, S. Yoo and D. Yu (2018). “KBase: The United States Department of Energy Systems Biology Knowledgebase.” Nature Biotechnology 36: 566. 10.1038/nbt.4163 https://www.nature.com/articles/nbt.4163#supplementary-information

Aziz, R. K., D. Bartels, A. A. Best, M. DeJongh, T. Disz, R. A. Edwards, K. Formsma, S. Gerdes, E. M. Glass, M. Kubal, F. Meyer, G. J. Olsen, R. Olson, A. L. Osterman, R. A. Overbeek, L. K. McNeil, D. Paarmann, T. Paczian, B. Parrello, G. D. Pusch, C. Reich, R. Stevens, O. Vassieva, V. Vonstein, A. Wilke and O. Zagnitko (2008). “The RAST Server: rapid annotations using subsystems technology.” BMC Genomics 9: 75. 10.1186/1471-2164-9-75

Baev, M. V., N. L. Schklyar, L. V. Chistoserdova, A. Y. Chistoserdov, B. M. Polanuer, Y. D. Tsygankov and V. E. Sterkin (1992). “Growth of the obligate methylotroph Methylobacillus flagellatum under stationary and nonstationary conditions during continuous cultivation.” Biotechnology and Bioengineering 39(6): 688–695. 10.1002/bit.260390614

Barber, R. D., M. A. Rott and T. J. Donohue (1996). “Characterization of a glutathione-dependent formaldehyde dehydrogenase from Rhodobacter sphaeroides.” Journal of Bacteriology 178(5): 1386–1393. doi:10.1128/jb.178.5.1386-1393.1996

Baumann, L., P. Baumann, M. Mandel and R. D. Allen (1972). “Taxonomy of Aerobic Marine Eubacteria.” Journal of Bacteriology 110(1): 402–429. doi:10.1128/jb.110.1.402-429.1972

Baumann, P., L. Baumann, R. Bowditch and B. Beaman (1984). “Taxonomy of Alteromonas: A. nigrifaciens sp. nov., nom. rev.; A macleodii; and A. Haloplanktis.” International Journal of Systematic and Evolutionary Microbiology 34. 10.1099/00207713-34-2-145

Bernstein, D. B., S. Sulheim, E. Almaas and D. Segrè (2021). “Addressing uncertainty in genome-scale metabolic model reconstruction and analysis.” Genome Biology 22(1): 64. 10.1186/s13059-021-02289-z

Bertilsson, S., O. Berglund, M. J. Pullin and S. W. Chisholm (2005). “Release of dissolved organic matter by Prochlorococcus.” Vie Et Milieu-Life and Environment 55(3-4): 225–231.

Bianchi, D., T. S. Weber, R. Kiko and C. Deutsch (2018). “Global niche of marine anaerobic metabolisms expanded by particle microenvironments.” Nature Geoscience 11(4): 263–268. 10.1038/s41561-018-0081-0

Biller, S. J., P. M. Berube, D. Lindell and S. W. Chisholm (2014). “Prochlorococcus: the structure and function of collective diversity.” Nature Reviews Microbiology 13: 13. 10.1038/nrmicro3378

Biller, S. J., A. Coe, A.-B. Martin-Cuadrado and S. W. Chisholm (2015). “Draft Genome Sequence of Alteromonas macleodii Strain MIT1002, Isolated from an Enrichment Culture of the Marine Cyanobacterium *Prochlorococcus*.” Genome Announcements 3(4): 10.1128/genomea.00967-00915. doi:10.1128/genomea.00967-15

Braakman, R., M. J. Follows and S. W. Chisholm (2017). “Metabolic evolution and the self-organization of ecosystems.” Proceedings of the National Academy of Sciences 114(15): E3091–E3100. 10.1073/pnas.1619573114

Buels, R., E. Yao, C. M. Diesh, R. D. Hayes, M. Munoz-Torres, G. Helt, D. M. Goodstein, C. G. Elsik, S. E. Lewis, L. Stein and I. H. Holmes (2016). “JBrowse: a dynamic web platform for genome visualization and analysis.” Genome Biology 17(1): 66. 10.1186/s13059-016-0924-1

Burley, S. K., C. Bhikadiya, C. Bi, S. Bittrich, L. Chen, G. V. Crichlow, C. H. Christie, K. Dalenberg, L. Di Costanzo, J. M. Duarte, S. Dutta, Z. Feng, S. Ganesan, D. S. Goodsell, S. Ghosh, R. K. Green, V. Guranović, D. Guzenko, B. P. Hudson, C. L. Lawson, Y. Liang, R. Lowe, H. Namkoong, E. Peisach, I. Persikova, C. Randle, A. Rose, Y. Rose, A. Sali, J. Segura, M. Sekharan, C. Shao, Y. P. Tao, M. Voigt, J. D. Westbrook, J. Y. Young, C. Zardecki and M. Zhuravleva (2021). “RCSB Protein Data Bank: powerful new tools for exploring 3D structures of biological macromolecules for basic and applied research and education in fundamental biology, biomedicine, biotechnology, bioengineering and energy sciences.” Nucleic Acids Res 49(D1): D437–d451. 10.1093/nar/gkaa1038

Cao, J.-Y., Y.-Y. Wang, M.-N. Wu, Z.-Y. Kong, J.-H. Lin, T. Ling, S.-M. Xu, S.-N. Ma, L. Zhang, C.-X. Zhou, X.-J. Yan and J.-L. Xu (2021). “RNA-seq Insights Into the Impact of Alteromonas macleodii on Isochrysis galbana.” Frontiers in Microbiology 12. 10.3389/fmicb.2021.711998

Caspi, R., R. Billington, I. M. Keseler, A. Kothari, M. Krummenacker, P. E. Midford, W. K. Ong, S. Paley, P. Subhraveti and P. D. Karp (2020). “The MetaCyc database of metabolic pathways and enzymes - a 2019 update.” Nucleic Acids Res 48(D1): D445–d453. 10.1093/nar/gkz862

Cecchini, G., I. Schröder, R. P. Gunsalus and E. Maklashina (2002). “Succinate dehydrogenase and fumarate reductase from Escherichia coli.” Biochimica et Biophysica Acta (BBA) - Bioenergetics 1553(1): 140–157. 10.1016/S0005-2728(01)00238-9

Concórdio-Reis, P., V. D. Alves, X. Moppert, J. Guézennec, F. Freitas and M. A. M. Reis (2021). “Characterization and Biotechnological Potential of Extracellular Polysaccharides Synthesized by Alteromonas Strains Isolated from French Polynesia Marine Environments.” Marine Drugs 19(9): 522.

Consortium, T. U. (2022). “UniProt: the Universal Protein Knowledgebase in 2023.” Nucleic Acids Research 51(D1): D523–D531. 10.1093/nar/gkac1052

Datta, M. S., A. A. Almada, M. F. Baumgartner, T. J. Mincer, A. M. Tarrant and M. F. Polz (2018). “Inter-individual variability in copepod microbiomes reveals bacterial networks linked to host physiology.” The ISME Journal 12(9): 2103–2113. 10.1038/s41396-018-0182-1

Diner, R. E., S. M. Schwenck, J. P. McCrow, H. Zheng and A. E. Allen (2016). “Genetic Manipulation of Competition for Nitrate between Heterotrophic Bacteria and Diatoms.” Frontiers in Microbiology 7(880). 10.3389/fmicb.2016.00880

Dixon, J. L., S. Sargeant, P. D. Nightingale and J. Colin Murrell (2013). “Gradients in microbial methanol uptake: productive coastal upwelling waters to oligotrophic gyres in the Atlantic Ocean.” The ISME Journal 7(3): 568–580. 10.1038/ismej.2012.130

Drula, E., M. L. Garron, S. Dogan, V. Lombard, B. Henrissat and N. Terrapon (2022). “The carbohydrate-active enzyme database: functions and literature.” 50(D1): D571–d577. 10.1093/nar/gkab1045

Dunn, N. A., D. R. Unni, C. Diesh, M. Munoz-Torres, N. L. Harris, E. Yao, H. Rasche, I. H. Holmes, C. G. Elsik and S. E. Lewis (2019). “Apollo: Democratizing genome annotation.” PLOS Computational Biology 15(2): e1006790. 10.1371/journal.pcbi.1006790

Elsik, C. G., K. C. Worley, L. Zhang, N. V. Milshina, H. Jiang, J. T. Reese, K. L. Childs, A. Venkatraman, C. M. Dickens, G. M. Weinstock and R. A. Gibbs (2006). “Community annotation: Procedures, protocols, and supporting tools.” Genome Research 16(11): 1329–1333. 10.1101/gr.5580606

Fadeev, E., F. De Pascale, A. Vezzi, S. Hübner, D. Aharonovich and D. Sher (2016). “Why close a bacterial genome? The plasmid of Alteromonas macleodii HOT1A3 is a vector for inter-specific transfer of a flexible genomic island.” Frontiers in Microbiology 7. 10.3389/fmicb.2016.00248

Fang, X., C. J. Lloyd and B. O. Palsson (2020). “Reconstructing organisms in silico: genome-scale models and their emerging applications.” Nature Reviews Microbiology 18(12): 731–743. 10.1038/s41579-020-00440-4

Finkel, Z. V., M. J. Follows, J. D. Liefer, C. M. Brown, I. Benner and A. J. Irwin (2016). “Phylogenetic Diversity in the Macromolecular Composition of Microalgae.” PLOS ONE 11(5): e0155977. 10.1371/journal.pone.0155977

Forchielli, E., D. Sher and D. Segrè (2022). “Metabolic Phenotyping of Marine Heterotrophs on Refactored Media Reveals Diverse Metabolic Adaptations and Lifestyle Strategies.” mSystems 0(0): e00070–00022. doi:10.1128/msystems.00070-22

Geider, R. J. and J. La Roche (2002). “Redfield revisited: variability of C: N: P in marine microalgae and its biochemical basis.” European Journal of Phycology 37(1): 1–17. Doi 10.1017/S0967026201003456

Gerlach, N. (2017). Biochemical and structural analysis of marine polysaccharide lyases. MSc, Universität Bremen.

Ghatak, S., Z. A. King, A. Sastry and B. O. Palsson (2019). “The y-ome defines the 35% of Escherichia coli genes that lack experimental evidence of function.” Nucleic Acids Res 47(5): 2446–2454. 10.1093/nar/gkz030

Givati, S., X. Yang, D. Sher and E. Rahav (2023). “Testing the growth rate and translation compensation hypotheses in marine bacterioplankton.” Environmental Microbiology **n/a**(n/a). 10.1111/1462-2920.16346

Green, M. L. and P. D. Karp (2004). “A Bayesian method for identifying missing enzymes in predicted metabolic pathway databases.” BMC Bioinformatics 5: 76. 10.1186/1471-2105-5-76

Harms, N., J. Ras, W. N. Reijnders, R. J. v. Spanning and A. H. Stouthamer (1996). “S- formylglutathione hydrolase of Paracoccus denitrificans is homologous to human esterase D: a universal pathway for formaldehyde detoxification?” Journal of Bacteriology 178(21): 6296–6299. doi:10.1128/jb.178.21.6296-6299.1996

Helland, S., B. F. Terjesen and L. Berg (2003). “Free amino acid and protein content in the planktonic copepod Temora longicornis compared to Artemia franciscana.” Aquaculture 215(1): 213–228. 10.1016/S0044-8486(02)00043-1

Hennon, G. M. M., J. J. Morris, S. T. Haley, E. R. Zinser, A. R. Durrant, E. Entwistle, T. Dokland and S. T. Dyhrman (2017). “The impact of elevated CO2 on Prochlorococcus and microbial interactions with ‘helper’ bacterium Alteromonas.” The Isme Journal 12: 520. 10.1038/ismej.2017.189 https://www.nature.com/articles/ismej2017189#supplementary-information

Henríquez-Castillo, C., A. M. Plominsky, S. Ramírez-Flandes, A. D. Bertagnolli, F. J. Stewart and O. Ulloa (2022). “Metaomics unveils the contribution of Alteromonas bacteria to carbon cycling in marine oxygen minimum zones.” Frontiers in Marine Science 9. 10.3389/fmars.2022.993667

Heyer, H. and W. E. Krumbein (1991). “Excretion of fermentation products in dark and anaerobically incubated cyanobacteria.” Archives of Microbiology 155(3): 284–287. 10.1007/BF00252213

Hobbs, J. K., S. M. Lee, M. Robb, F. Hof, C. Barr, K. T. Abe, J.-H. Hehemann, R. McLean, D. W. Abbott and A. B. Boraston (2016). “KdgF, the missing link in the microbial metabolism of uronate sugars from pectin and alginate.” Proceedings of the National Academy of Sciences 113(22): 6188–6193. doi:10.1073/pnas.1524214113

Hosmani, P. S., T. Shippy, S. Miller, J. B. Benoit, M. Munoz-Torres, M. Flores-Gonzalez, L. A. Mueller, H. Wiersma-Koch, T. D’Elia, S. J. Brown and S. Saha (2019). “A quick guide for student-driven community genome annotation.” PLOS Computational Biology 15(4): e1006682. 10.1371/journal.pcbi.1006682

Hou, S., M. López-Pérez, U. Pfreundt, N. Belkin, K. Stüber, B. Huettel, R. Reinhardt, I. Berman-Frank, F. Rodriguez-Valera and W. R. Hess (2018). “Benefit from decline: the primary transcriptome of Alteromonas macleodii str. Te101 during Trichodesmium demise.” The ISME Journal 12(4): 981–996. 10.1038/s41396-017-0034-4

Huang, L., Y. Zhang, X. Du, R. An and X. Liang (2022). “Escherichia coli Can Eat DNA as an Excellent Nitrogen Source to Grow Quickly.” Frontiers in Microbiology 13. 10.3389/fmicb.2022.894849

Ivars-Martinez, E., A.-B. Martin-Cuadrado, G. D’Auria, A. Mira, S. Ferriera, J. Johnson, R. Friedman and F. Rodriguez-Valera (2008). “Comparative genomics of two ecotypes of the marine planktonic copiotroph Alteromonas macleodii suggests alternative lifestyles associated with different kinds of particulate organic matter.” Isme J 2(12): 1194–1212.

Jung, H., T. Ventura, J. S. Chung, W.-J. Kim, B.-H. Nam, H. J. Kong, Y.-O. Kim, M.-S. Jeon and S.-i. Eyun (2020). “Twelve quick steps for genome assembly and annotation in the classroom.” PLOS Computational Biology 16(11): e1008325. 10.1371/journal.pcbi.1008325

Kallen, R. G. and W. P. Jencks (1966). “The Mechanism of the Condensation of Formaldehyde with Tetrahydrofolic Acid.” Journal of Biological Chemistry 241(24): 5851–5863. 10.1016/S0021-9258(18)96350-7

Kanehisa, M. and S. Goto (2000). “KEGG: Kyoto Encyclopedia of Genes and Genomes.” Nucleic Acids Research 28(1): 27–30. 10.1093/nar/28.1.27

Karlsen, E., C. Schulz and E. Almaas (2018). “Automated generation of genome-scale metabolic draft reconstructions based on KEGG.” BMC Bioinformatics 19(1): 467. 10.1186/s12859-018-2472-z

Karp, P. D., R. Billington, R. Caspi, C. A. Fulcher, M. Latendresse, A. Kothari, I. M. Keseler, M. Krummenacker, P. E. Midford, Q. Ong, W. K. Ong, S. M. Paley and P. Subhraveti (2019). “The BioCyc collection of microbial genomes and metabolic pathways.” Brief Bioinform 20(4): 1085–1093. 10.1093/bib/bbx085

Karp, P. D., P. E. Midford, R. Billington, A. Kothari, M. Krummenacker, M. Latendresse, W. K. Ong, P. Subhraveti, R. Caspi, C. Fulcher, I. M. Keseler and S. M. Paley (2021). “Pathway Tools version 23.0 update: software for pathway/genome informatics and systems biology.” Brief Bioinform 22(1): 109–126. 10.1093/bib/bbz104

Karp, P. D., S. Paley, R. Caspi, A. Kothari, M. Krummenacker, P. E. Midford, L. R. Moore, P. Subhraveti, S. Gama-Castro, V. H. Tierrafria, P. Lara, L. Muñiz-Rascado, C. Bonavides-Martinez, A. Santos-Zavaleta, A. Mackie, G. Sun, T. A. Ahn-Horst, H. Choi, M. W. Covert, J. Collado-Vides and I. Paulsen (2023). “The EcoCyc Database (2023).” EcoSal Plus 0(0): eesp-0002-2023. doi:10.1128/ecosalplus.esp-0002-2023

Katoh, K. and D. M. Standley (2013). “MAFFT Multiple Sequence Alignment Software Version 7: Improvements in Performance and Usability.” Molecular Biology and Evolution 30(4): 772–780. 10.1093/molbev/mst010

Kazakov, A. E., D. A. Rodionov, E. Alm, A. P. Arkin, I. Dubchak and M. S. Gelfand (2009). “Comparative Genomics of Regulation of Fatty Acid and Branched-Chain Amino Acid Utilization in Proteobacteria.” Journal of Bacteriology 191(1): 52–64. doi:10.1128/jb.01175-08

Kearney, S. M., E. Thomas, A. Coe and S. W. Chisholm (2021). “Microbial diversity of co-occurring heterotrophs in cultures of marine picocyanobacteria.” Environmental Microbiome 16(1): 1. 10.1186/s40793-020-00370-x

Keltjens, J. T., A. Pol, J. Reimann and H. J. Op den Camp (2014). “PQQ-dependent methanol dehydrogenases: rare-earth elements make a difference.” Appl Microbiol Biotechnol 98(14): 6163–6183. 10.1007/s00253-014-5766-8

Koch, H., A. Dürwald, T. Schweder, B. Noriega-Ortega, S. Vidal-Melgosa, J.-H. Hehemann, T. Dittmar, H. M. Freese, D. Becher, M. Simon and M. Wietz (2019). “Biphasic cellular adaptations and ecological implications of Alteromonas macleodii degrading a mixture of algal polysaccharides.” The ISME Journal 13(1): 92–103. 10.1038/s41396-018-0252-4

Lee, C., S. G. Wakeham and J. I. Hedges (2000). “Composition and flux of particulate amino acids and chloropigments in equatorial Pacific seawater and sediments.” Deep Sea Research Part I: Oceanographic Research Papers 47(8): 1535–1568. 10.1016/S0967-0637(99)00116-8

Lee, M. D., N. G. Walworth, E. L. McParland, F.-X. Fu, T. J. Mincer, N. M. Levine, D. A. Hutchins and E. A. Webb (2017). “The Trichodesmium consortium: conserved heterotrophic co-occurrence and genomic signatures of potential interactions.” The ISME Journal 11(8): 1813–1824. 10.1038/ismej.2017.49

Lee, T. J., I. Paulsen and P. Karp (2008). “Annotation-based inference of transporter function.” Bioinformatics 24(13): i259–267. 10.1093/bioinformatics/btn180

Lemos, M. L., A. E. Toranzo and J. L. Barja (1985). “Modified Medium for the Oxidation-Fermentation Test in the Identification of Marine Bacteria.” Applied and Environmental Microbiology 49(6): 1541–1543. doi:10.1128/aem.49.6.1541-1543.1985

Li, W., K. R. O’Neill, D. H. Haft, M. DiCuccio, V. Chetvernin, A. Badretdin, G. Coulouris, F. Chitsaz, Myra K. Derbyshire, A. S. Durkin, N. R. Gonzales, M. Gwadz, Christopher J. Lanczycki, J. S. Song, N. Thanki, J. Wang, Roxanne A. Yamashita, M. Yang, C. Zheng, A. Marchler-Bauer and F. Thibaud-Nissen (2020). “RefSeq: expanding the Prokaryotic Genome Annotation Pipeline reach with protein family model curation.” Nucleic Acids Research 49(D1): D1020–D1028. 10.1093/nar/gkaa1105

Lidbury, I., J. C. Murrell and Y. Chen (2014). “Trimethylamine *N*-oxide metabolism by abundant marine heterotrophic bacteria.” Proceedings of the National Academy of Sciences 111(7): 2710–2715. doi:10.1073/pnas.1317834111

Liu, X., Y. Dong, J. Zhang, A. Zhang, L. Wang and L. Feng (2009). “Two novel metal-independent long-chain alkyl alcohol dehydrogenases from Geobacillus thermodenitrificans NG80-2.” Microbiology 155(6): 2078–2085. 10.1099/mic.0.027201-0

Liu, Y., L. Sargent, W. Leung, S. C. R. Elgin and J. Goecks (2019). “G-OnRamp: a Galaxy-based platform for collaborative annotation of eukaryotic genomes.” Bioinformatics 35(21): 4422–4423. 10.1093/bioinformatics/btz309

Lopez-Perez, M., A. Gonzaga, A. B. Martin-Cuadrado, O. Onyshchenko, A. Ghavidel, R. Ghai and F. Rodriguez-Valera (2012). “Genomes of surface isolates of Alteromonas macleodii: the life of a widespread marine opportunistic copiotroph.” Sci Rep 2: 696. 10.1038/srep00696

López-Pérez, M., N. Ramon-Marco and F. Rodriguez-Valera (2017). “Networking in microbes: conjugative elements and plasmids in the genus Alteromonas.” BMC Genomics 18(1): 36. 10.1186/s12864-016-3461-0

López-Pérez, M. and F. Rodriguez-Valera (2016). “Pangenome evolution in the marine bacterium Alteromonas.” Genome Biology and Evolution. 10.1093/gbe/evw098

Maia, L. B., L. Fonseca, I. Moura and J. J. G. Moura (2016). “Reduction of Carbon Dioxide by a Molybdenum-Containing Formate Dehydrogenase: A Kinetic and Mechanistic Study.” Journal of the American Chemical Society 138(28): 8834–8846. 10.1021/jacs.6b03941

Manck, L. E., J. Park, B. J. Tully, A. M. Poire, R. M. Bundy, C. L. Dupont and K. A. Barbeau (2022). “Petrobactin, a siderophore produced by Alteromonas, mediates community iron acquisition in the global ocean.” The ISME Journal 16(2): 358–369. 10.1038/s41396-021-01065-y

Markowitz, V. M., K. Mavromatis, N. N. Ivanova, I. M. Chen, K. Chu and N. C. Kyrpides (2009). “IMG ER: a system for microbial genome annotation expert review and curation.” Bioinformatics 25(17): 2271–2278. btp393 [pii] 10.1093/bioinformatics/btp393

Marx, C. J., L. Chistoserdova and M. E. Lidstrom (2003a). “Formaldehyde-Detoxifying Role of theTetrahydromethanopterin-Linked Pathway in *Methylobacteriumextorquens*AM1.” Journal of Bacteriology 185(24): 7160–7168. doi:10.1128/jb.185.23.7160-7168.2003

Marx, C. J., M. Laukel, J. A. Vorholt and M. E. Lidstrom (2003b). “Purification of the Formate-Tetrahydrofolate Ligasefrom *Methylobacterium extorquens* AM1 and Demonstrationof Its Requirement for MethylotrophicGrowth.” Journal of Bacteriology 185(24): 7169–7175. doi:10.1128/jb.185.24.7169-7175.2003

McCarren, J., J. W. Becker, D. J. Repeta, Y. Shi, C. R. Young, R. R. Malmstrom, S. W. Chisholm and E. F. DeLong (2010). “Microbial community transcriptomes reveal microbes and metabolic pathways associated with dissolved organic matter turnover in the sea.” Proc Natl Acad Sci U S A 107(38): 16420–16427.

Mehta, A., C. Sidhu, A. K. Pinnaka and A. Roy Choudhury (2014). “Extracellular polysaccharide production by a novel osmotolerant marine strain of Alteromonas macleodii and its application towards biomineralization of silver.” PLoS One 9(6): e98798. 10.1371/journal.pone.0098798

Mestre, M., E. Borrull, M. M. Sala and J. M. Gasol (2017). “Patterns of bacterial diversity in the marine planktonic particulate matter continuum.” The ISME Journal 11(4): 999–1010. 10.1038/ismej.2016.166

Métris, A., P. Sudhakar, D. Fazekas, A. Demeter, E. Ari, M. Olbei, P. Branchu, R. A. Kingsley, J. Baranyi and T. Korcsmáros (2017). “SalmoNet, an integrated network of ten Salmonella enterica strains reveals common and distinct pathways to host adaptation.” npj Systems Biology and Applications 3(1): 31. 10.1038/s41540-017-0034-z

Mincer, T. J. and A. C. Aicher (2016). “Methanol Production by a Broad Phylogenetic Array of Marine Phytoplankton.” PLOS ONE 11(3): e0150820. 10.1371/journal.pone.0150820

Moisander, P. H., K. M. Shoemaker, M. C. Daley, E. McCliment, J. Larkum and M. A. Altabet (2018). “Copepod-Associated Gammaproteobacteria Respire Nitrate in the Open Ocean Surface Layers.” Frontiers in Microbiology 9. 10.3389/fmicb.2018.02390

Morris, J. J., R. Kirkegaard, M. J. Szul, Z. I. Johnson and E. R. Zinser (2008). ”Facilitation of robust growth of Prochlorococcus colonies and dilute liquid cultures by “helper” heterotrophic bacteria.” Appl Environ Microbiol 74(14): 4530–4534.

Mueller, L. A., P. Zhang and S. Y. Rhee (2003). “AraCyc: a biochemical pathway database for Arabidopsis.” Plant Physiol 132(2): 453–460. 10.1104/pp.102.017236

Müller, J. E. N., F. Meyer, B. Litsanov, P. Kiefer and J. A. Vorholt (2015). “Core pathways operating during methylotrophy of Bacillus methanolicus MGA3 and induction of a bacillithiol-dependent detoxification pathway upon formaldehyde stress.” Molecular Microbiology 98(6): 1089–1100. 10.1111/mmi.13200

Neumann, A. M., J. P. Balmonte, M. Berger, H.-A. Giebel, C. Arnosti, S. Voget, M. Simon, T. Brinkhoff and M. Wietz (2015). “Different utilization of alginate and other algal polysaccharides by marine Alteromonas macleodii ecotypes.” Environmental Microbiology 17(10): 3857–3868. 10.1111/1462-2920.12862

Paley, S., P. E. O’Maille, D. Weaver and P. D. Karp (2016). “Pathway collages: personalized multi-pathway diagrams.” BMC Bioinformatics 17(1): 529. 10.1186/s12859-016-1382-1

Pedler, B. E., L. I. Aluwihare and F. Azam (2014). “Single bacterial strain capable of significant contribution to carbon cycling in the surface ocean.” Proceedings of the National Academy of Sciences 111(20): 7202–7207. 10.1073/pnas.1401887111

Pedreira, T., C. Elfmann and J. Stülke (2021). “The current state of SubtiWiki, the database for the model organism Bacillus subtilis.” Nucleic Acids Research 50(D1): D875–D882. 10.1093/nar/gkab943

Raguénès, G., M. A. Cambon-Bonavita, J. F. Lohier, C. Boisset and J. Guezennec (2003). “A Novel, Highly Viscous Polysaccharide Excreted by an Alteromonas Isolated from a Deep-Sea Hydrothermal Vent Shrimp.” Current Microbiology 46(6): 0448–0452. 10.1007/s00284-002-3922-3

Ramsey, J., B. McIntosh, D. Renfro, S. A. Aleksander, S. LaBonte, C. Ross, A. E. Zweifel, N. Liles, S. Farrar, J. J. Gill, I. Erill, S. Ades, T. Z. Berardini, J. A. Bennett, S. Brady, R. Britton, S. Carbon, S. M. Caruso, D. Clements, R. Dalia, M. Defelice, E. L. Doyle, I. Friedberg, S. M. R. Gurney, L. Hughes, A. Johnson, J. M. Kowalski, D. Li, R. C. Lovering, T. L. Mans, F. McCarthy, S. D. Moore, R. Murphy, T. D. Paustian, S. Perdue, C. N. Peterson, B. M. Prüß, M. S. Saha, R. R. Sheehy, J. T. Tansey, L. Temple, A. W. Thorman, S. Trevino, A. C. Vollmer, V. Walbot, J. Willey, D. A. Siegele and J. C. Hu (2021). “Crowdsourcing biocuration: The Community Assessment of Community Annotation with Ontologies (CACAO).” PLOS Computational Biology 17(10): e1009463. 10.1371/journal.pcbi.1009463

Reintjes, G., C. Arnosti, B. Fuchs and R. Amann (2019). “Selfish, sharing and scavenging bacteria in the Atlantic Ocean: a biogeographical study of bacterial substrate utilisation.” The ISME Journal 13(5): 1119–1132. 10.1038/s41396-018-0326-3

Ro, Y. T., H. I. Lee, E. J. Kim, J. H. Koo, E. Kim and Y. M. Kim (2003). “Purification, characterization, and physiological response of a catalase-peroxidase in Mycobacterium sp. strain JC1 DSM 3803 grown on methanol.” FEMS Microbiology Letters 226(2): 397–403. 10.1016/s0378-1097(03)00644-x

Romero, P., J. Wagg, M. L. Green, D. Kaiser, M. Krummenacker and P. D. Karp (2005). “Computational prediction of human metabolic pathways from the complete human genome.” Genome Biol 6(1): R2. 10.1186/gb-2004-6-1-r2

Roth Rosenberg, D., M. Haber, J. Goldford, M. Lalzar, D. Aharonovich, A. Al-Ashhab, Y. Lehahn, D. Segrè, L. Steindler and D. Sher (2021). “Particle-associated and free-living bacterial communities in an oligotrophic sea are affected by different environmental factors.” Environmental Microbiology n/a(n/a). 10.1111/1462-2920.15611

Satish Kumar, V., M. S. Dasika and C. D. Maranas (2007). “Optimization based automated curation of metabolic reconstructions.” BMC Bioinformatics 8(1): 212. 10.1186/1471-2105-8-212

Schulz-Mirbach, H., A. Müller, T. Wu, P. Pfister, S. Aslan, L. Schada von Borzyskowski, T. J. Erb, A. Bar-Even and S. N. Lindner (2022). “On the flexibility of the cellular amination network in E coli.” eLife 11: e77492. 10.7554/eLife.77492

Seaver, S. M. D., F. Liu, Q. Zhang, J. Jeffryes, J. P. Faria, J. N. Edirisinghe, M. Mundy, N. Chia, E. Noor, M. E. Beber, A. A. Best, M. DeJongh, J. A. Kimbrel, P. D’Haeseleer, S. R. McCorkle, J. R. Bolton, E. Pearson, S. Canon, E. M. Wood-Charlson, R. W. Cottingham, A. P. Arkin and C. S. Henry (2021). “The ModelSEED Biochemistry Database for the integration of metabolic annotations and the reconstruction, comparison and analysis of metabolic models for plants, fungi and microbes.” Nucleic Acids Res 49(D1): D575–d588. 10.1093/nar/gkaa746

Seemann, T. (2014). “Prokka: rapid prokaryotic genome annotation.” Bioinformatics 30(14): 2068–2069. 10.1093/bioinformatics/btu153

Shane, B. (1989). Folylpolyglutamate Synthesis and Role in the Regulation of One-Carbon Metabolism. Vitamins & Hormones. G. D. Aurbach and D. B. McCormick, Academic Press. 45: 263–335. 10.1016/S0083-6729(08)60397-0

Sheik, A. R., C. P. Brussaard, G. Lavik, P. Lam, N. Musat, A. Krupke, S. Littmann, M. Strous and M. M. Kuypers (2014). “Responses of the coastal bacterial community to viral infection of the algae Phaeocystis globosa.” ISME J 8(1): 212–225. 10.1038/ismej.2013.135

Shibl, A. A., A. Isaac, M. A. Ochsenkühn, A. Cárdenas, C. Fei, G. Behringer, M. Arnoux, N. Drou, M. P. Santos, K. C. Gunsalus, C. R. Voolstra and S. A. Amin (2020). “Diatom modulation of select bacteria through use of two unique secondary metabolites.” Proceedings of the National Academy of Sciences 117(44): 27445–27455. doi:10.1073/pnas.2012088117

Sosa, O. A., J. R. Casey and D. M. Karl (2019). “Methylphosphonate Oxidation in *Prochlorococcus* Strain MIT9301 Supports Phosphate Acquisition, Formate Excretion, and Carbon Assimilation into Purines.” Applied and Environmental Microbiology 85(13): e00289–00219. 10.1128/aem.00289-19

Srivastava, A., D. E. M. Saavedra, B. Thomson, J. A. L. García, Z. Zhao, W. M. Patrick, G. J. Herndl and F. Baltar (2021). “Enzyme promiscuity in natural environments: alkaline phosphatase in the ocean.” The ISME Journal 15(11): 3375–3383. 10.1038/s41396-021-01013-w

Stein, L. (2001). “Genome annotation: from sequence to biology.” Nature Reviews Genetics 2(7): 493–503. 10.1038/35080529

Sterck, L., K. Billiau, T. Abeel, P. Rouzé and Y. Van de Peer (2012). “ORCAE: online resource for community annotation of eukaryotes.” Nature Methods 9(11): 1041–1041. 10.1038/nmeth.2242

Sun, J., L. Steindler, J. C. Thrash, K. H. Halsey, D. P. Smith, A. E. Carter, Z. C. Landry and S. J. Giovannoni (2011). “One Carbon Metabolism in SAR11 Pelagic Marine Bacteria.” PLOS ONE 6(8): e23973. 10.1371/journal.pone.0023973

Tarazona-Janampa, U. I., A. D. Cembella, M. C. Pelayo-Zárate, S. Pajares, L. M. Márquez-Valdelamar, Y. B. Okolodkov, J. Tebben, B. Krock and L. M. Durán-Riveroll (2020). “Associated Bacteria and Their Effects on Growth and Toxigenicity of the Dinoflagellate Prorocentrum lima Species Complex From Epibenthic Substrates Along Mexican Coasts.” Frontiers in Marine Science 7. 10.3389/fmars.2020.00569

Tinta, T., Z. Zhao, B. Bayer and G. J. Herndl (2023). “Jellyfish detritus supports niche partitioning and metabolic interactions among pelagic marine bacteria.” Microbiome 11(1): 156. 10.1186/s40168-023-01598-8

Vries, G. E. d., N. Arfman, P. Terpstra and L. Dijkhuizen (1992). “Cloning, expression, and sequence analysis of the Bacillus methanolicus C1 methanol dehydrogenase gene.” Journal of Bacteriology 174(16): 5346–5353. doi:10.1128/jb.174.16.5346-5353.1992

Weissberg, O., D. Aharonovich and D. Sher (2023). “Phototroph-heterotroph interactions during growth and long-term starvation across Prochlorococcus and Alteromonas diversity.” The ISME Journal 17(2): 227–237. 10.1038/s41396-022-01330-8

Wietz, M., M. López-Pérez, D. Sher, S. J. Biller and F. Rodriguez-Valera (2022). “Microbe Profile: Alteromonas macleodii - a widespread, fast-responding, ‘interactive’ marine bacterium.” Microbiology (Reading) 168(11). 10.1099/mic.0.001236

Zhang, H., T. Yohe, L. Huang, S. Entwistle, P. Wu, Z. Yang, P. K. Busk, Y. Xu and Y. Yin (2018). “dbCAN2: a meta server for automated carbohydrate-active enzyme annotation.” Nucleic Acids Res 46(W1): W95–w101. 10.1093/nar/gky418

Zhao, Z., F. Baltar and G. J. Herndl (2020). “Linking extracellular enzymes to phylogeny indicates a predominantly particle-associated lifestyle of deep-sea prokaryotes.” Science Advances 6(16): eaaz4354. doi:10.1126/sciadv.aaz4354

Zoccarato, L., D. Sher, T. Miki, D. Segrè and H.-P. Grossart (2022). “A comparative whole-genome approach identifies bacterial traits for marine microbial interactions.” Communications Biology 5(1): 276. 10.1038/s42003-022-03184-4

